# Single cell analysis uncovers striking cellular heterogeneity of lung infiltrating Tregs during eosinophilic vs. neutrophilic allergic airway inflammation

**DOI:** 10.1101/2023.09.23.559152

**Authors:** Supinya Iamsawat, Rongzhen Yu, Sohee Kim, Kevin Qiu, Jaehyuk Choi, Booki Min

**Affiliations:** Department of Microbiology and Immunology, Northwestern University Feinberg School of Medicine, Chicago, IL 60611; Department of Inflammation and Immunity, Lerner Research Institute, Cleveland Clinic Foundation, Cleveland, OH 44195; Department of Dermatology, Northwestern University Feinberg School of Medicine, Chicago, IL 60611; Department of Biochemistry and Molecular Genetics, Northwestern University Feinberg School of Medicine, Chicago, IL 60611

**Author notes:** These authors contributed equally (Supinya Iamsawat, Rongzhen Yu). Corresponding author: Booki Min, Department of Microbiology and Immunology, Northwestern University Feinberg School of Medicine, 310 E Superior St., Morton 6-626, Chicago, IL 60611. EMAIL).

## Abstract

Allergic airway inflammation is a chronic inflammatory condition resulting from uncontrolled immune responses to environmental antigens. While it is well-established that allergic immune responses exhibit a high degree of diversity, driven by primary effector cell types like eosinophils, neutrophils, or CD4 T cells with distinct effector signatures, the exact mechanisms responsible for such pathogenesis remain largely elusive. Foxp3^+^ regulatory T cells (Treg) is an essential immune regulator during chronic inflammation including allergic airway inflammation. Emerging evidence suggests that Tregs infiltrating inflamed tissues exhibit distinct phenotypes dependent on the specific tissue sites and can display significant heterogeneity and tissue residency. Whether diverse allergic inflammatory responses in the lung influences infiltrating Treg heterogeneity or Treg lung residency has not previously been explored. We employed an unbiased single-cell RNAseq approach to investigate lung-infiltrating Tregs in models of eosinophilic and neutrophilic airway inflammation models, in which Tregs are critical regulators of inflammation. We found that lung-infiltrating Tregs are highly heterogeneous and that Tregs displaying lung resident phenotypes are significantly different depending on the types of inflammation. Tregs expression of ST2, a receptor for alarmin cytokine IL-33, was predominantly induced by eosinophilic inflammation and by tissue residency. However, Treg-specific ST2 deficiency did not affect the development of eosinophilic allergic inflammation nor the generation of lung resident Tregs. These results uncover a striking heterogeneity among Tregs infiltrating the lungs during allergic airway inflammation. The results also indicate that varying types of inflammation may give rise to phenotypically distinct lung resident Tregs, underscoring a novel mechanism by which inflammatory cues may shape the composition of infiltrating Tregs, allowing them to regulate inflammatory responses through tissue-adapted mechanisms.

## Introduction

CD4 T cell immunity mediating allergic inflammatory responses in the lung comes in different flavors. Conventional allergic responses are often associated with type 2 immunity displaying signature cytokines, IL-5 and IL-13. On the other hand, proinflammatory T cells with Th1 and Th17 type signature cytokines are also found associated with certain types of allergic inflammation, particularly severe steroid-resistant form of inflammation (1, 2). In our prior study, we demonstrated two distinct experimental models of allergic airway inflammation, each eliciting specific types of inflammatory responses based on the choice of adjuvants used during sensitization: alum adjuvant-induced eosinophilic inflammation mediated by effector CD4 T cells with type 2 cytokine profiles and CFA adjuvant-induced neutrophilic inflammation mediated by CD4 T cells displaying type 1/17 cytokine profiles (3). The mechanisms responsible for these inflammatory endotypes remain largely unknown.

Foxp3^+^ regulatory T cells (Tregs) play an indispensable role in regulating inflammatory responses. During allergic inflammation, Tregs are essential to regulate immune responses as well as to promote tissue repair and airway remodeling process (4, 5). Indeed, it was shown that Treg depletion during sensitization was sufficient to aggravate allergic airway inflammation (6). We previously reported that Treg depletion during antigen challenge similarly exacerbates allergic inflammation, regardless of the adjuvants used (3). From flow cytometric analysis we noticed some differences in Treg marker expression, such as ICOS and Nrp1, although other Treg markers including CTLA4 and GITR were expressed at comparable levels (3).

The goal of this study is to unbiasedly examine lung infiltrating Tregs during steroid-sensitive eosinophilic and steroid-resistant neutrophilic airway inflammation to test the hypothesis that lung infiltrating Tregs under these conditions display qualitatively different gene profiles. Through nanostring and single cell RNA sequencing analyses we discovered that the population of Tregs infiltrating the lung tissue is notably diverse, including highly suppressive phenotype resembling non-lymphoid tissue-like effector Treg populations and highly proliferative central memory cell-like Treg populations. We also employed an in vivo labeling approach to separately analyze lung resident and intravascular Tregs and found that lung resident Tregs in eosinophilic inflammation were highly enriched in ST2^+^ Tregs, where IL-33 production is pronounced. However, deletion of ST2 expression specifically in Tregs did not affect Treg-mediated control of eosinophilic airway inflammation. Our data not only suggest the complexity of Treg-dependent regulation of allergic airway inflammation but also demonstrate an intricate interplay between tissue-derived inflammatory factors and tissue resident Tregs.

## Materials and Methods

### Animals

C57BL/6 Foxp3^GFP^-knockin mice were previously reported (7). C57BL/6 Foxp3^Cre^*Il1rl*^fl/fl^ mice were kindly provided by Dr. Diane Mathis (Harvard Medical School) (8). All the animal experiments were approved by the institutional animal care and use committee of Northwestern University. All the methods were performed in accordance with the relevant guidelines and regulations. All mice were bred in a specific pathogen free facility at Northwestern University Feinberg School of Medicine.

### Allergic airway inflammation

Eosinophilic airway inflammation was induced by two intraperitoneal injections of 5μg of cockroach antigen (CA, Greer Laboratory, Lenoir, NC) mixed in 100μl of alum adjuvant (aluminum hydroxide, Sigma, St. Louis, MO) at day 0 and 7. Starting on day 14, the sensitized mice were intranasally challenged daily with CA (5μg in 50μl saline) for 4 consecutive days. Animals were sacrificed 24 hours after the last CA challenge.

Neutrophilic airway inflammation was induced by subcutaneous injection of 5μg of CA emulsidied in 100μl of CFA containing 5mg/ml H37Ra (BD Difco, Franklin Lakes, NJ). Intranasal CA challenge was performed as described above. For in vivo labeling, the mice were intravenously injected with 1.5μg of PE-conjugated anti-CD4 (RM4-4) antibody in 100μl sterile saline and sacrificed 5 minutes after the injection.

### Flow cytometry

Bronchoalveolar lavage (BAL) fluid cells were collected for analysis. Lung tissue was minced and digested with collagenase and DNase, and a single cell suspension was obtained. Mononuclear cells were isolated using a Percoll gradient centrifugation. RBCs were then lysed using ACK lysis buffer. Cells were stained with anti-CD4 (RM4-5), Ly6G (1A8), SiglecF (1RNM44N), CTLA4 (UC10-4B9), OX40 (OX-86), Foxp3 (FJK-16s),

ICOS (C398.4A), PD1 (J43), and Ki67 (B56) antibodies. All the antibodies used were purchased from Biolegend (San Diego, CA), eBioscience (San Diego, CA), and BD PharMingen (San Jose, CA). For intracellular cytokine staining, cells were stimulated ex vivo with PMA (10ng/ml) and ionomycin (1μM) for 4 hours in the presence of 2μM monensin (Calbiochem, San Diego, CA) during the last 2 hours of stimulation. Cells were acquired using a FACS Symphony A1 flow cytometer and analyzed using a FlowJo software (Treestar, Ashland, OR).

### Single cell RNAseq and data analysis

Tregs were FACS sorted using a FACS Melody cell sorter (BD Bioscience), and then captured for library generation and sequencing using 10X Chromium single cell capture chip according to the manufacturers protocol. The single-cell RNAseq libraries were prepared according to manufacturer’s Chromium Single Cell 3ʹ protocol (10X Genomics, document # CG000204). In brief, >5,000 FACS sorted Tregs were targeted using the 10x Genomics Chromium Controller for cDNA synthesis and barcoding. cDNA quality and quantity of each sample were assessed using a Bioanalyzer High Sensitivity DNA assay. cDNA was used to begin fragmentation, followed by end-repair, adapter ligation, and sample indexing. Sample libraries were then assessed for proper construction using the same Bioanalyzer assay. Constructed libraries were pooled and quantified using a Quantabio Q cycler, and then denatured and sequenced on an Illumina Novaseq 6000 high-throughput sequencing platform with 28 cycles for the forward read and 91 cycles for the reverse read. Each cell was sequenced to a minimum of 20,000 reads per cell per manufacturer’s recommendations. The 10X results were subjected to processing through the Cell Ranger pipeline version 3.1.0 to generate Fastq files. Fastq files were subsequently aligned using the Cell Ranger Count v3.1.0 pipeline within the 10X Genomics cloud analysis platform (https://www.10xgenomics.com/products/cloud-analysis), employing the Single Cell 3’ Gene Expression library type. Filtered gene expression matrices obtained from the cloud analysis were imported into R (v4.1.3) and processed using the Seurat package (v4.3.0.1) (9). Cells were filtered based on the criteria of nFeature_RNA > 500 and < 6000, and percent.mito < 9.15 and normalization was performed using the SCTransform function. Pseudotime analysis and Branch Expression Analysis Modeling (BEAM) were executed using the monocle package (v2.22.0) (10–12). Enriched pathways of genes identified from the BEAM plot were subjected to analysis using g:Profiler (Database version: 29/03/2023) (13). Single cell RNAseq data was deposited to GEO (GSE243653).

### Quantitative real time PCR analysis

RNA was extracted from the total lung tissues using a GeneJET RNA isolation kit (Thermo Fisher, Waltham, MA) and cDNA was synthesized using M-MLV reverse transcriptase (Promega, Madison, WI). Real time quantitative PCR analysis was performed using a QuantStudio 3 real time PCR system (Applied Biosystems, Waltham, MA) using a Radiant qPCR mastermix (Alkali Scientific, Fort Lauderdale, FL). The data were normalized by housekeeping *Gapdh* gene and the relative expression was calculated. Taqman primers used are: *Il4* (Mm00445259_m1), *Il13* (Mm00434204_m1), *Il17a* (Mm00439618_m1), *Ifng* (Mm01168134_m1), *Il1rl1* (Mm00516117_m1), and *Il33* (Mm00505403_m1).

### Nanostring analysis

Nanostring analysis to measure gene expression was previously reported (14). In brief, 100ng of total RNA was hybridized to the Mouse Immunology Panel overnight and then processed on a GEN2 analyst system using the high-sensitivity protocol and high-resolution data capture. Raw counts were normalized using a nSolver 2.5 software. Background counts were removed by subtracting the mean plus 2 SDs of the negative controls, and lane specific differences in hybridization and binding intensity were corrected using the geometric mean of the positive controls. Gene expression was normalized to the geometric mean of three reference genes (*Oaz1*, *G6pd*, and *Gapdh*).

### Statistical analysis

Statistical significance was determined by the Mann-Whitney test using a Prism software (GraphPad, San Diego, CA). p<0.05 was considered statistically significant.

## Results

### Gene expression profiles in lung Tregs during eosinophilic vs. neutrophilic airway inflammation

Asthmatic inflammation in the lung is highly heterogeneous, and the cellular and molecular mechanisms underlying such different endotypes remain largely unknown. Experimentally, using different adjuvants for antigen sensitization could elicit different types of effector immunity. We previously reported two different regimens to sensitize animals with cockroach antigen (CA) in either alum or CFA adjuvant. Following intranasal CA challenge, the recipients mount distinct airway inflammation depending on the adjuvants used (3). For instance, broncho alveolar lavage (BAL) cells from alum-induced inflammation featured predominantly eosinophils with minor neutrophils, whereas neutrophilic proportions were dramatically increased in BAL cells from CFA-induced inflammation (Fig 1A). Effector T cell immunity mediating the inflammation is also distinct. Alum-induced inflammation is typically associated with CD4 T cells with Th2 type signature, whereas CFA-induced inflammation is characterized by Th1/Th17 type T cell immunity (3). In support, cytokine gene expression of the lung tissue measured by qPCR analysis clearly supported Th2 and Th1/Th17 immunity during alum- and CFA-induced airway inflammation, respectively (Fig 1B). Interestingly, alarmin *Il33* and its receptor *Il1rl1* expression was clearly increased in alum-induced inflammation model (Fig 1B). In this study, alum- and CFA-induced inflammation will be used to refer eosinophilic and neutrophilic airway inflammation, respectively.

**Figure 1.**
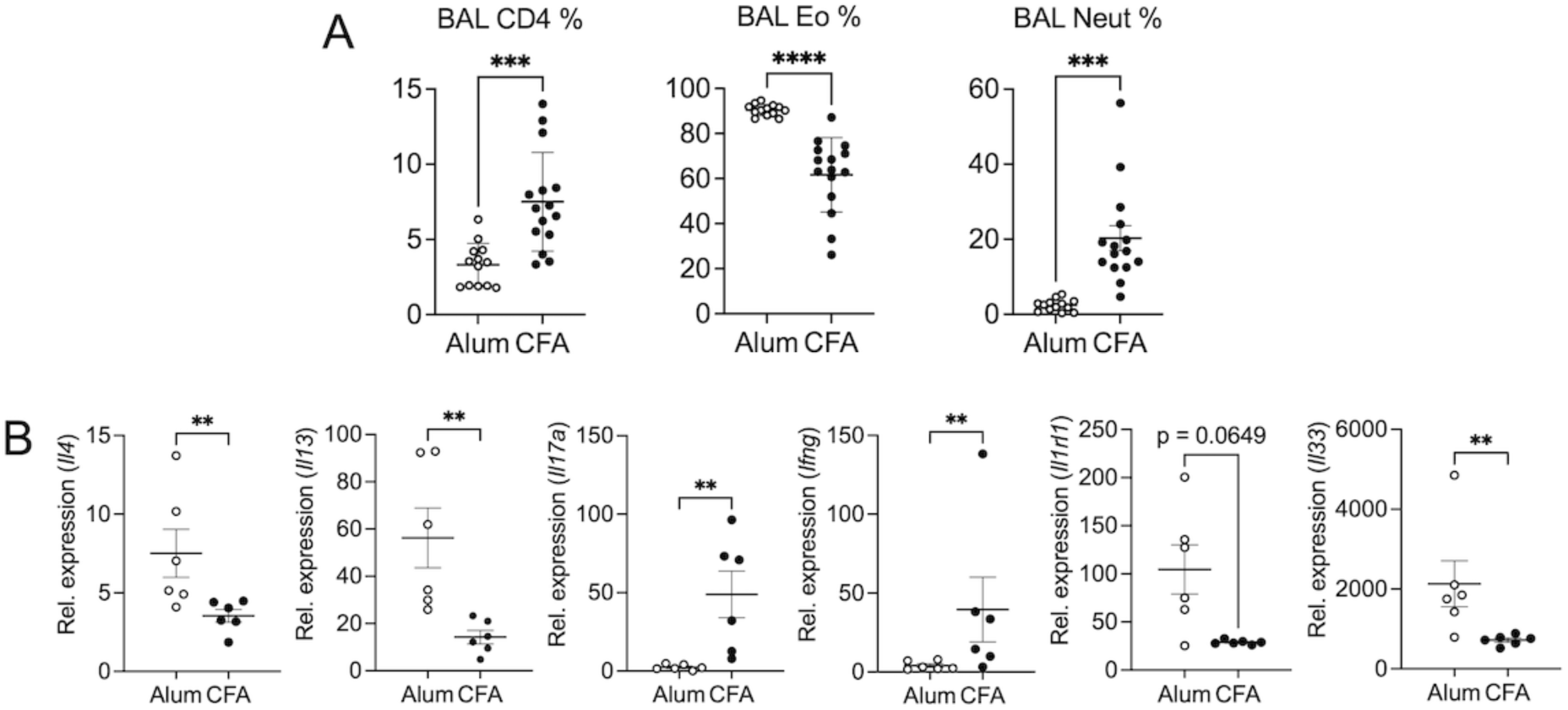
**Alum-and CFA-induced airway inflammation models.** Mice were sensitized with CA in either alum or CFA adjuvant, followed by intranasal CA challenge as described in Materials and Methods. (A) Upon sacrifice, BAL cells were harvested. Proportion of CD4 T cells, eosinophils, and neutrophils was determined. (B) Expression of the indicated genes in the lung tissue was determined by qPCR analysis. Each symbol represents individually tested animal. **, p<0.01; ***, p<0.001; ****, p<0.0001.

We previously reported that Tregs are essential regulators of both alum- and CFA-induced allergic inflammation and that they may exploit different mechanisms to control inflammatory responses (3). To closely investigate Tregs during different allergic inflammatory responses in the lung, lung infiltrating Tregs were FACS sorted and examined for gene expression profiles by a Nanostring analysis. Fig 2A and 2B showed genes differentially expressed in Tregs from each inflammatory condition. Genes associated with alum-induced inflammation include *Il17rb* (receptor for IL-25), *Il10*, *Il1rl1* (receptor for IL-33), and *Tnfaip3*. Metascape analysis of those differentially expressed genes identified pathways highly enriched in Tregs from alum-induced inflammation, and these include positive regulation of cytokine production and chronic inflammatory response (Fig 2C). On the other hand, Treg genes associated with CFA-induced inflammation were Th1/Th17-related proinflammatory genes, such as *Tbx21*, *Il17a*, *Il12rb1*, *Il12rb2*, *Cxcr3*, and *Irf1* (Fig 2A and 2B). We also tested pathways enriched in these Tregs, and they are cytokine-cytokine receptor interaction and cytokine mediated signaling pathway (Fig 2C). Of note, Foxp3 expression level was comparable between the groups (Fig 2B). Therefore, analyzing different gene expression of the lung Tregs from these experimental models corroborates the earlier hypothesis that Tregs may exert regulatory functions through diverse mechanisms. Because tissue infiltrating Tregs could be highly heterogeneous, we hypothesize that such distinct mechanisms may be mediated by different Treg subsets.

**Figure 2.**
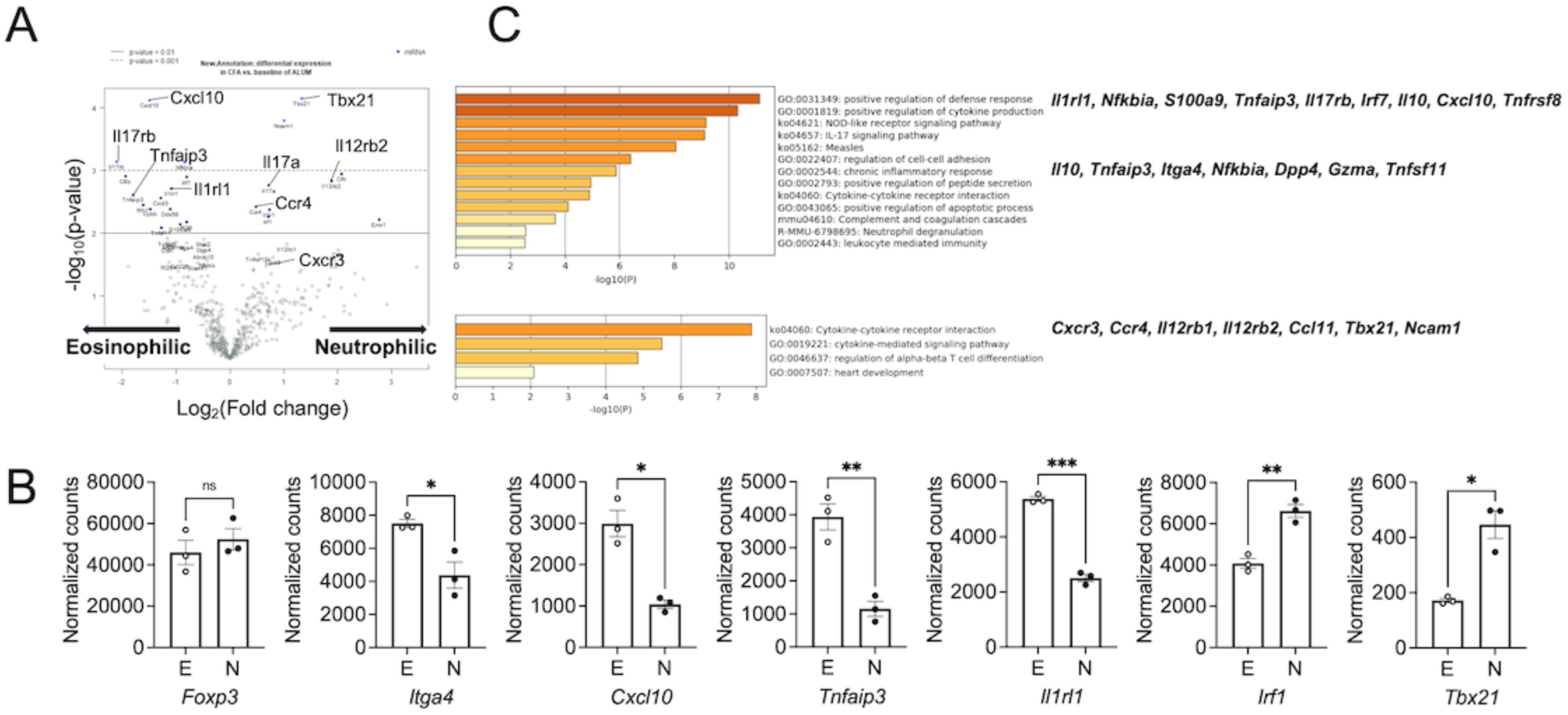
**Nanostring analysis of lung Tregs** Lung infiltrating Tregs were isolated from mice with alum-induced eosinophilic or CFA-induced neutrophilic airway inflammation as described in the Materials and Methods. (A) Volcano plots represent differentially expressed genes. (B) Normalized counts of the indicated genes. (C) Differentially expressed genes were subjected to Metascape analysis to identify key pathways highly enriched in each inflammation condition. *, p<0.05; **, p<0.01; ***, p<0.001.

### scRNAseq analysis of lung infiltrating Tregs during eosinophilic and neutrophilic airway inflammation

The Treg heterogeneity within different tissue sites has well been appreciated, and tissue-derived factors such as antigens and cytokines may contribute to the heterogeneity (15–17). To test if inflammatory conditions play a role in such Treg heterogeneity, we conducted a single cell RNAseq experiment using Tregs isolated from the lung tissues of Foxp3^GFP^ mice undergoing alum- and CFA-induced airway inflammation. Uniform Manifold Approximation and Projection (UMAP) analysis demonstrated substantial heterogeneity of those Tregs infiltrating the lung tissues (Fig 3A). Three clusters (clusters 0, 1, 2) expressed genes associated with non-lymphoid tissue (NLT) Treg signatures. Cluster 0 resembled NLT-like effector Tregs based on their high expression of *Maf*, *Cxcr6*, *Pim1*, and *Itgae* (gene encoding CD103) (18).

**Figure 3.**
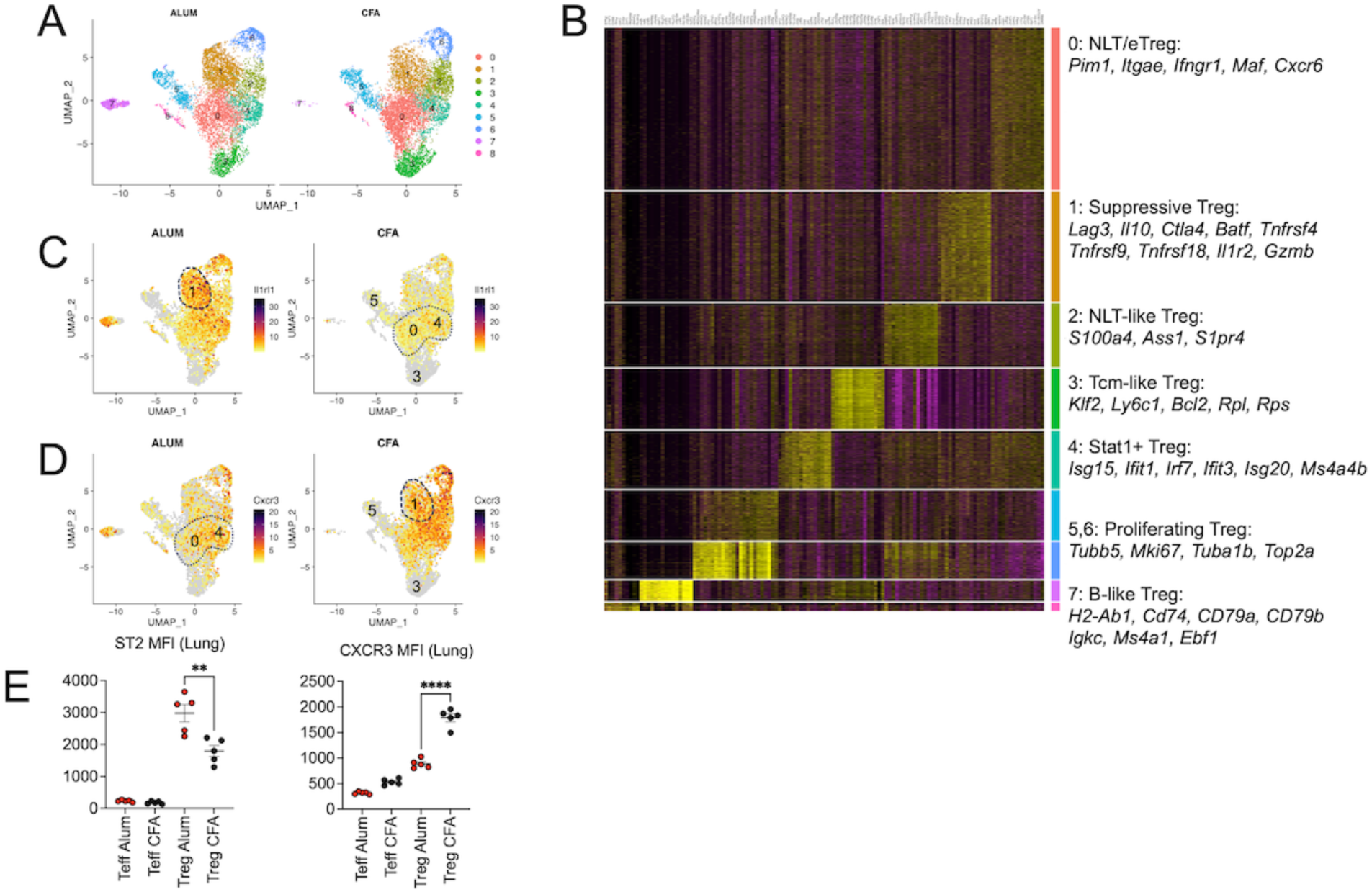
**Heterogeneity of lung infiltrating Tregs** Lung infiltrating Tregs from alum-induced and CFA-induced airway inflammation were subjected to single cell RNAseq analysis. (A) UMAP plots of Tregs from each condition. (B) Heatmap plots showing top differentially expressed genes from each cluster. (C and D) Feature plots showing *Il1rl1* (C) and *Cxcr3* (D) expressing Tregs. (E) Foxp3^GFP^ mice induced for alum-induced or CFA-induced airway inflammation were examined for ST2 and CXCR3 expression in lung infiltrating effector CD4 (Teff) or Tregs. **, p<0.01; ****, p<0.0001.

Cluster 1 was characterized by genes associated with Treg suppressive functions, including *Lag3*, *Il10*, *Ctla4*, and TNF receptor superfamilies such as *Tnfrsf4* (OX40), *Tnfrsf9* (4-1BB), and *Tnfrsf18* (GITR). In addition to those NLT and potentially suppressive Treg subsets, a distinct NLT-like Treg subset that expresses different gene sets were also identified. Cluster 2 Tregs expressed *S100a4*, *S100a10*, *Ass1*, *Lsp1*, and *S1pr4*, which were previously found highly enriched in NLT-like Tregs (19). We also found Tregs expressing central memory T cell-like genes (cluster 3), and they are *Klf2*, *Ly6c*, *Bcl2*, and many ribosome subunit genes (20–22). Treg cluster featuring IFN-responsive genes (cluster 4), including *Isg15*, *Ifit1*, *Irf7*, *Ifit3*, and *Isg20*, has previously been identified and referred to as Stat1^+^ Tregs (19). In addition, two closely related clusters (5 and 6) of Tregs characterized by genes associated with proliferation were noted. They expressed tubulin genes (*Tubb5* and *Tuba1b*), topoisomerase (*Top2a*), and *Mki67*. Notably, their expression appears to be more pronounced in the cluster 6 (Fig 3B). Interestingly, we also observed a B cell-like Treg subset (cluster 7) that highly expressed B cell genes such as *Cd74*, *Cd79*, *Igkc*, *Ms4a1*, and *Ebf1* gene. Interestingly, this cluster was predominantly found in Tregs from alum-induced inflammation (Fig 3A). Since ST2 (encoded by the *Il1rl1* gene) and CXCR3 expression was especially greater in Tregs from alum-and CFA-induced inflammation models (Fig 2A), respectively, we next examined Tregs expressing the *Il1rl1* and *Cxcr3* gene. Feature plots showed that *Il1rl1*^+^ Tregs were found in NLT-like, Stat1^+^, and proliferating Treg clusters. In particular, suppressive Treg (cluster 1) highly expressed the *Il1rl1* gene, and more importantly, they were more evident in Tregs from alum-induced inflammation (Fig 3C). On the other hand, *Cxcr3* expression was most pronounced in the similar clusters of Tregs from CFA-induced inflammation (Fig 3D). Likewise, Cxcr3 expression of the suppressive Treg cluster 1 was greater in CFA-induced than that in alum-induced inflammation (Fig 3D). Flow cytometry measurement of ST2 and CXCR3 expression in Tregs further confirmed Treg expression of these molecules (Fig 3E). The data also validated greater expression of ST2 and CXCR3 in Tregs from alum- and CFA-induced inflammation, respectively (Fig 3E).

### Re-clustering analysis of Tregs expressing suppressive phenotypes

We noticed significant differences in the proportions of each Treg cluster between the two inflammation models. For instance, Tregs from CFA-induced inflammation exhibited a higher abundance of clusters 0 and 2, while Tregs from alum-induced inflammation displayed larger clusters 1 and 7 (Supp Fig S1). We were particularly interested in the cluster 1, which highly expressed genes associated with Treg suppressive functions. Since *Il1rl1*^+^ and *Cxcr3*^+^ Tregs are especially enriched in the cluster 1 in alum- and CFA-induced inflammation, respectively (Fig 3C and 3D), we wanted to test if Tregs that belong to cluster 1 might be distinct between the two models. We thus re-clustered suppressive Treg cluster 1. UMAP analysis from the re-clustered population showed a striking difference between the two inflammation models (Fig 4A). From the UMAP plot, the new cluster 0 was predominantly found in Tregs from CFA-induced inflammation, whereas the new clusters 1-3 were primarily observed in Tregs from alum-induced inflammation (Fig 4A). The new Treg cluster 0 expressed *Cxcr3* and *Ms4a6b* (Fig 4B). It was previously reported that a subset of Tregs express CXCR3 to acquire trafficking pattern of pathogenic Th1 cells, which then support Treg mobilization like Th1 cells and Treg suppression of pathogenic Th1 responses at sites of inflammation (23). In support, Treg expression of Cxcr3 was mainly found in the clusters from CFA-induced inflammation group (Fig 4C). Ms4a6b is a member of MS4A genes, whose expression is highly Treg specific, and upregulated by TGFβ or Foxp3 (24). Moreover, it was shown that MS4a6b interacts with GITR in Tregs and amplifies signals from TCR and GITR (24). On the other hand, *Il1rl1*, *Gata3*, and *Fgl2* were highly expressed in clusters 1-3, which were mainly found in Tregs from alum-induced inflammation (Fig 4A). The IL-33/ST2 pathway was recently shown to be critical to control Treg regulatory functions (25–27), although the opposite impact of IL-33 on Treg functions has also been reported (28). ST2 expression identifies highly activated suppressive Tregs preferentially located in non-lymphoid tissues (29, 30). Indeed, ST2 expression of Tregs was mostly found in the clusters from alum-induced inflammation model (Fig 4C).

**Figure 4.**
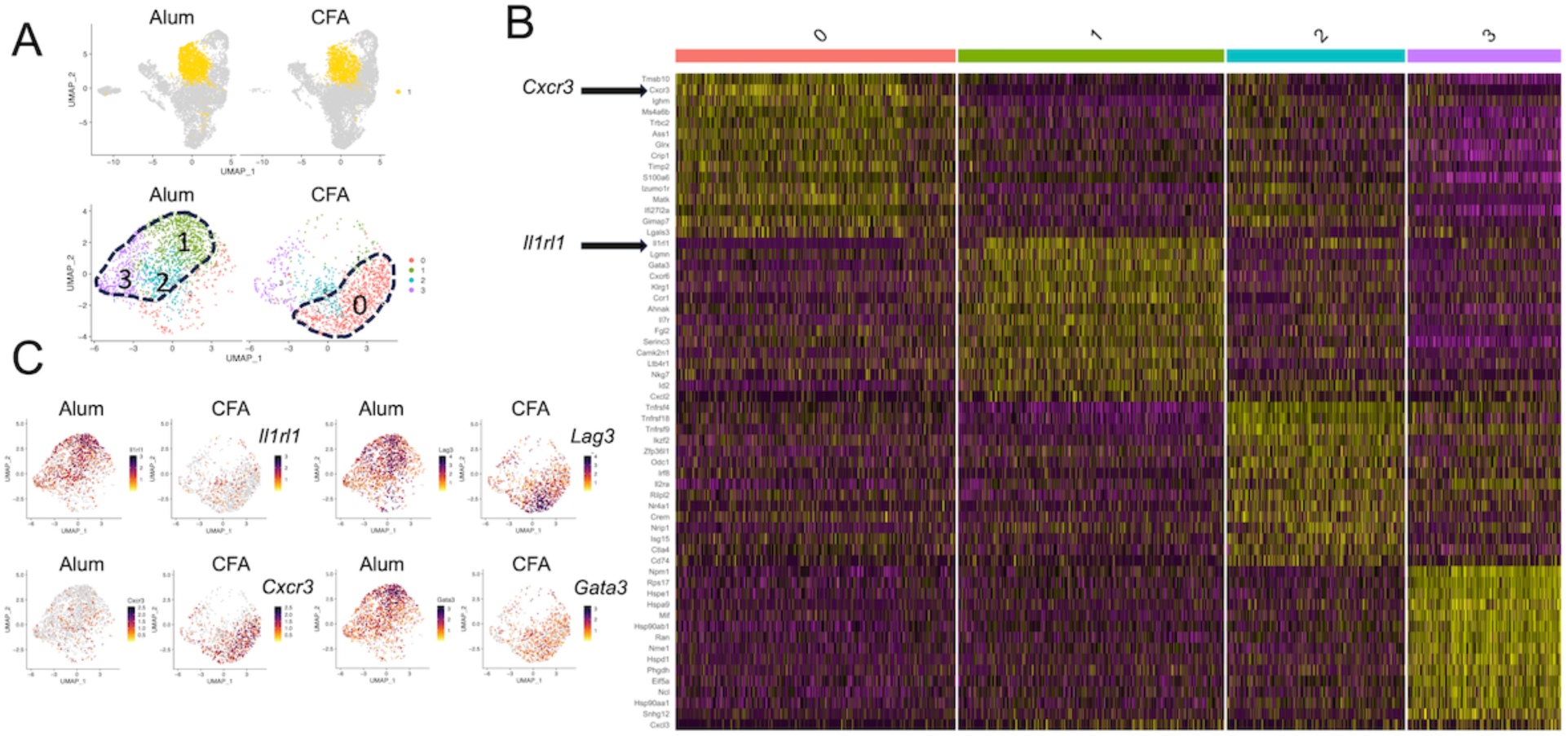
**Re-clustering analysis of Tregs that express suppressive markers.** (A) Cluster #1 of the Figure 2, Tregs with suppressive phenotypes, was re-clustered. The new UMAP plot shows distinct Treg clusters between the inflammation models. (B) Heatmap plots showing top differentially expressed genes from each cluster. (C) Feature plots showing *Il1rl1*, *Lag3*, *Cxcr3*, and *Gata3* expression.

Similar re-cluster analysis was conducted for cluster 0 (from Fig 3B) that express NLT-like effector Treg markers. As shown in Supp Fig S2, re-clustering showed substantial heterogeneity between the groups, although the differences were not as evident as cluster 1 (Supp Fig S2A). Of note, new clusters 2 and 3 were evident in Tregs from CFA-induced inflammation, whereas clusters 4 and 5 was abundant in Tregs from alum-induced inflammation (Supp Fig S2A). Interestingly, *Il1rl1* and *Cxcr3* expressing Tregs from this population also appeared to be distinct, as *Il1rl1*^+^ Tregs were mostly found in clusters 4 and 5 from alum-induced inflammation and in clusters 2 and 3 from CFA-induced inflammation (Supp Fig S2B). Likewise, *Cxcr3*^+^ Tregs predominantly localized in the clusters 2 and 3 in CFA-induced inflammation, and their expression in Tregs from alum-induced inflammation was sparsely found in cluster 4 and 5 (Supp Fig S2B). Collectively, these results demonstrate that Tregs infiltrating the lung tissues are highly heterogeneous and that some Treg subsets, especially with suppressive phenotypes, were found different depending on the types of inflammation.

### The hierarchy of lung infiltrating Tregs differentiation during allergic inflammation

To delineate the hierarchy and transition relationships between different Treg subsets (NLT, NLT-like, Tcm-like, etc), we next performed a pseudotime trajectory analysis. The trajectories from the analysis predicted a branch with one major split, which divided the cells into three states: pre-branch, cell fate 1, and cell fate 2 (Fig 5A). Tregs from the ‘pre-branch’ group were associated with both proliferative and suppressive clusters (Fig 5B). Tregs transitioning into cell fate 1 displayed traits of central memory T cell-like subsets (Fig 5B). Expression of proliferation markers, Mki67 and Top2a, was also high at the pre-branch proliferating subsets and gradually declined in suppressive subsets (Supp Fig S2C). Tregs then regained proliferative ability once they differentiated into central memory T-like subsets during which Tregs greatly increase Bcl2 and Klf2 expression (Fig 5C and Supp Fig S2C). In support, pathways related to cytosolic ribosomes, translation, and biosynthetic processes were significantly enriched during this fate determination (Fig 5C). On the other hand, ‘pre-branch’ Tregs became NLT-like effector Treg subsets during cell fate 2 determination (Fig 5B). Importantly, while expression of key Treg markers associated with suppressive ability, such as Ctla4, Lag3, Il10, Tnfrsf4 (OX40), and Tnfrsf9 (4-1BB), was gradually diminished during central memory T-like Treg differentiation, their expression was maintained during cell fate 2 differentiation (Fig 5C). Moreover, pathways related to immune processes, cytokine stimulation, and T cell activation, were highly enriched in this process (Fig 5C).

**Figure 5.**
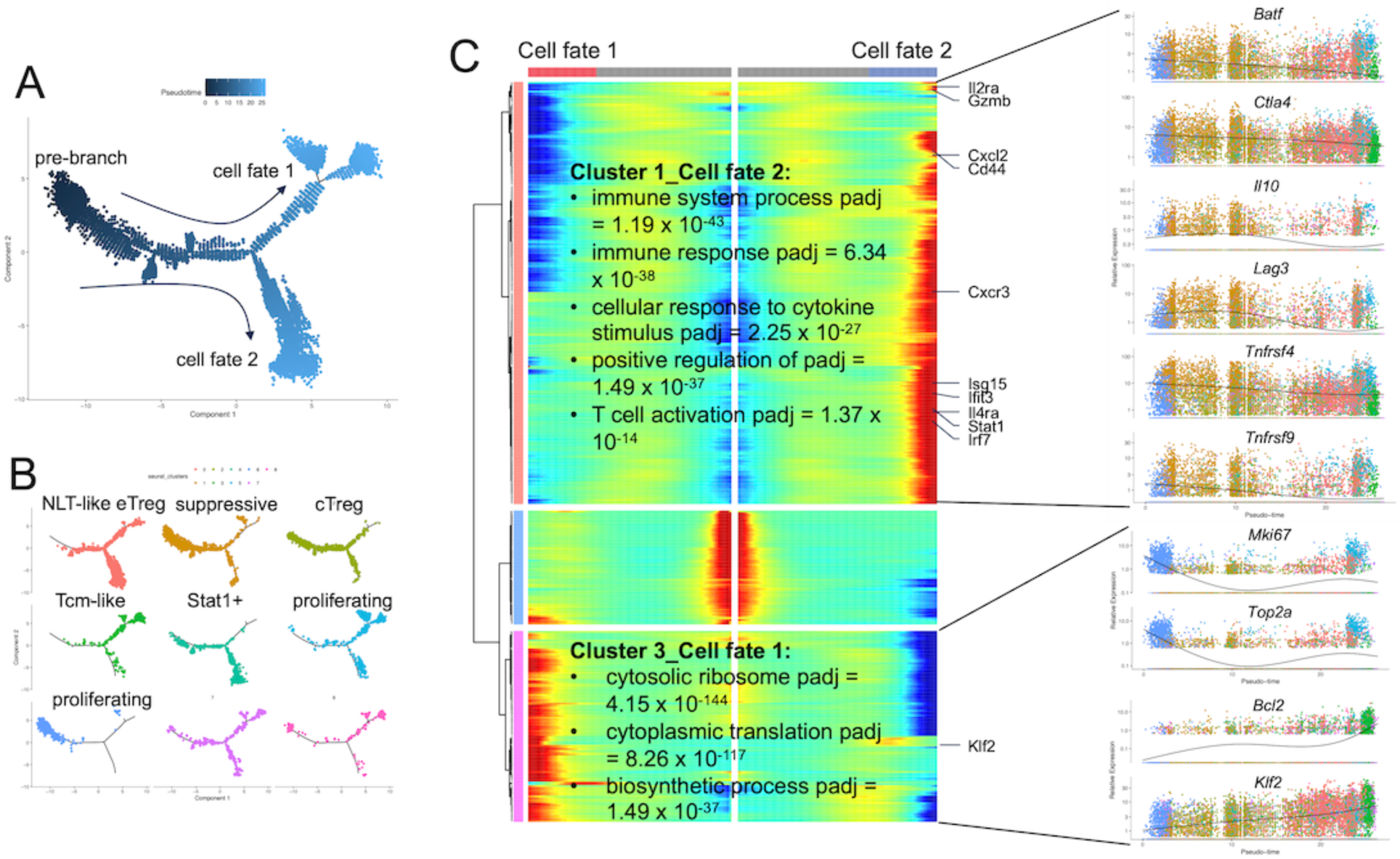
**Pseudotime trajectory analysis of Tregs** (A) Monocle pseudotime trajectory of lung Tregs. (B) Each cluster was plotted by pseudotime. (C) Differentially expressed genes for each cell fate pathway were plotted in a BEAM plot. Each column represents a cell, and each row represents a gene. Key pathways enriched in each cell fate were analyzed. Pseudotime ordered single cell expression of representative genes are shown.

### Lung resident Tregs are ST2^high^ cells during alum-induced airway inflammation

It was previously reported that lung resident and circulating memory T cells express distinct functions in supporting allergic airway inflammation (31). We thus sought to compare circulating and lung resident Tregs during alum- and CFA-induced airway inflammation. In vivo labeling of intravascular cells with fluorochrome conjugated antibodies was performed to distinguish lung resident cells from cells in the vasculature as previously reported (32). Intravascular (i.v. anti-CD4 positive) and lung resident (i.v. anti-CD4 negative) T cells were separately analyzed (Fig 6A). The proportions of total CD4 T cells and of Tregs were comparable between alum- and CFA-induced inflammations (Fig 6B). The proportions of lung resident Foxp3^-^ effector CD4 T cells were significantly higher in CFA-induced inflammation, while lung resident Treg proportions were similar (Fig 6C). We then compared phenotypes of intravascular and lung resident effector CD4 T cells and Tregs and found that Foxp3 expression level was significantly higher in lung resident Tregs compared to that in intravascular Tregs (Fig 6D). Proliferation activity determined by Ki67 staining showed that Tregs were generally more proliferative then effector T cells and that lung resident subsets were also more proliferative than intravascular subsets. Expression of other Treg associated molecules, namely OX40, ICOS, PD1, and intracellular CTLA4, followed similar expression pattern (Fig 6D). The sources of Tregs (i.e., alum- or CFA-induced inflammation) did not affect the expression patterns of these molecules (Fig 6D). However, we noticed that CD103 expression was significantly higher in lung resident effector T cells from CFA-induced inflammation (Fig 6E). CD103 expression of intravascular Tregs was comparable between the two models, although it was greater in lung resident Tregs from CFA-induced inflammation (Fig 6E). Interestingly, T cell expression of ST2 was predominantly observed from alum-induced inflammation model (Fig 6F). Moreover, Treg expression of ST2 was substantially higher, and more importantly, lung resident Tregs expressed even higher level of ST2 compared to that of intravascular Tregs (Fig 6F). Therefore, these results demonstrate that ST2 is a unique Treg marker associated with alum-induced airway inflammation.

**Figure 6.**
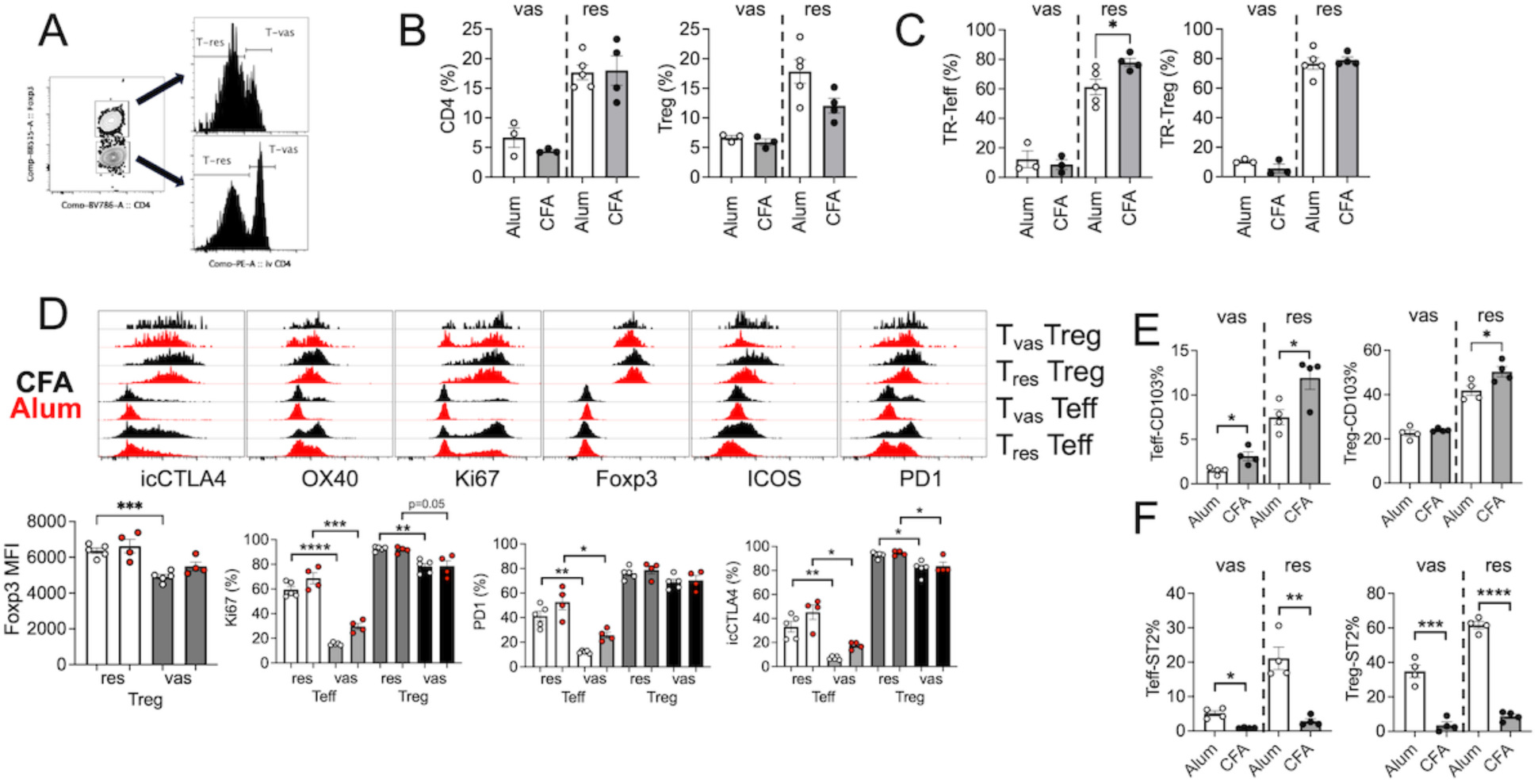
**Lung resident and intravascular Tregs in alum- and CFA-induced airway inflammation.** (A) In vivo anti-CD4 antibody labeling of lung cells. Lung resident (res) and intravascular (vas) effector CD4 and Tregs are shown. (B) Proportions of CD4 T cells and Tregs in the lung at the peak of the airway inflammation. (C) Lung resident and intravascular effector CD4 T cells and Tregs were examined. (D) Phenotypic analysis of lung resident and intravascular effector CD4 T cells and Tregs measuring intracellular CTLA4, OX40, Ki-67, Foxp3, ICOS, and PD-1 expression. (E and F) Proportion of CD103^+^ (E) and ST2^+^ (F) lung resident (res) and intravascular (vas) effector CD4 T cells (Teff) and Tregs (Treg) in alum- and CFA-induced airway inflammation. Each symbol represents individually tested animal. The data represent two independent experiments. *, p<0.05; **, p<0.01; ***, p<0.001; ****, p<0.0001.

### ST2 signaling in Tregs is dispensable for Treg-mediated immune regulation during alum-induced airway inflammation

The importance of the IL-33/ST2 axis in Treg function has been explored in several inflammation models, including parasite infection and autoimmunity (33–35). However, the role of IL-33 in Treg function during allergic inflammation remains controversial (25, 28). Based on the findings that ST2 is preferentially expressed in Tregs during alum-induced airway inflammation, we next investigated whether ST2 expression by Tregs is necessary for Tregs to control inflammation, especially alum-induced inflammation model. Treg-specific ST2 deficient (Treg^ΔST2^) animals were thus sensitized with alum adjuvant and induced for allergic inflammation. Treg-specific ST2 deficiency was validated by flow analysis (data not shown). Since lung resident subsets expressed significantly higher ST2 compared to intravascular cells (Fig 6F), we separately analyzed T cell subsets based on their location. The level of lung infiltrating Tregs was similar regardless of ST2 expression (Fig 7A). When compared for tissue-residency, we found that lung resident effector CD4 T cells were significantly reduced in Treg^ΔST2^ group, although lung resident Tregs remained comparable (Fig 7B). Proliferative activity of intravascular and lung resident effector CD4 and Tregs remained similar, suggesting that the lack of ST2 expression in Tregs did not affect the cellular proliferation in vivo (Fig 7C and 7D). Moreover, expression of PD1, OX40, and intracellular CTLA4 in the indicated T cell subsets was not statistically different (Fig 7C and 7D). When T cell expression of inflammatory cytokine IL-13 was measured, we found no differences in cytokine expressing CD4 T cells in the lung (Fig 7E). IL-33-responsive Tregs were previously shown to limit γδ T cell responses in the lung or in the central nervous system during autoimmune neuroinflammation (25, 35). However, we found no difference in IL-17-producing γδ T cells in Treg^ΔST2^ mice (Fig 7F). Therefore, our data suggest that, despite its prominent expression in lung-resident Tregs, IL-33 signaling plays a limited or negligible role in modulating inflammatory T cell responses and the tissue residency of Tregs in the lung.

**Figure 7.**
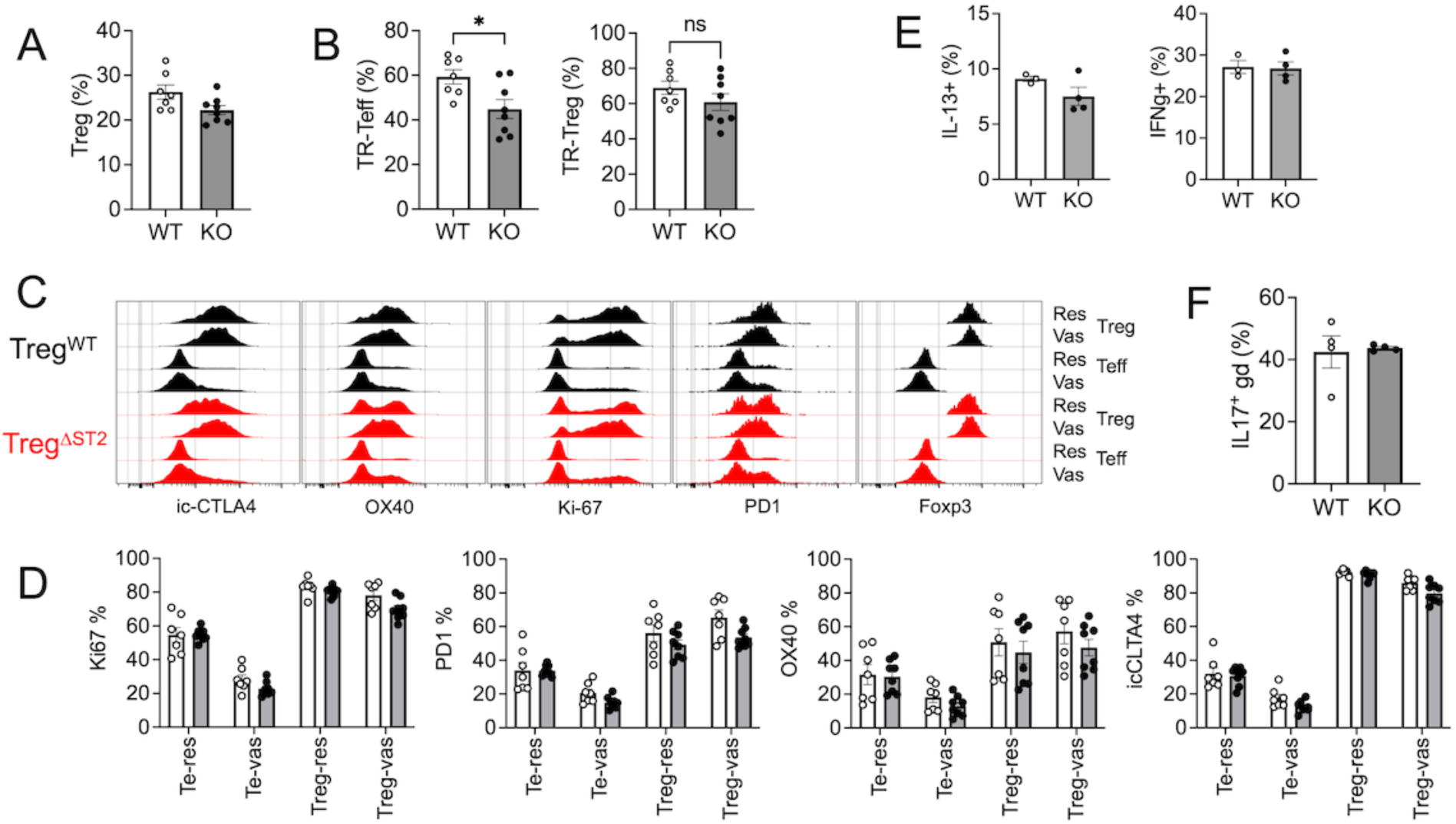
**Alum-induced eosinophilic airway inflammation in Treg specific ST2 deficient animals.** Groups of Treg specific ST2 deficient (Treg^ΔST2^) and littermate wild type control mice were induced for eosinophilic airway inflammation as described in Materials and Methods. (A) Lung infiltrating Tregs at the peak of the responses. (B) Proportion of lung resident effector CD4 T cells and Tregs. (C and D) Phenotypic analysis of lung resident and intravascular effector CD4 T cells and Tregs. (E) CD4 T cell expression of IL-13 and IFNγ was determined. (F) IL-17^+^ ψ8 T cells in the lung. Each symbol represents individually tested animal. The data shown are a representative of two independent experiments. *, p<0.05.

## Discussion

The pathogenesis of allergic inflammatory responses in the lung is intricate, mediated by multiple cell types that exert both positive and negative control over these processes. Effector CD4 T cells with Th2 signature are considered major effector cells in conventional allergic inflammation, while Th17 type effector CD4 T cells are thought to mediate non-conventional allergic inflammation that is often more severe and refractory to traditional intervention such as steroids (36–38). By utilizing different adjuvants during sensitization, we previously reported two distinct allergic airway inflammation models characterized by predominant infiltration of eosinophils and CD4 T cells producing IL-5 and IL-13 or by mixture of eosinophils and drastically elevated neutrophils along with CD4 T cells expressing Th1/Th17 type cytokines (3). While effector immunity mediating the inflammation may be distinct, we found that Foxp3^+^ Tregs are instrumental in limiting both types of inflammation, because Treg depletion exacerbated the inflammation regardless of the types of immune responses (3). Although we previously concluded that Tregs may utilize different mechanisms to control the inflammation, the precise nature of Treg-dependent regulation of different allergic inflammation remains largely unexplored.

Tregs are present within the non-lymphoid tissues, such as adipose tissue, muscle, and skin, and they are thought to play an important role in tissue homeostasis (39, 40). The inflamed lung tissues during allergic airway inflammation are heavily infiltrated with Tregs; however, the mechanisms through which they control lung inflammation has not thoroughly been examined. Single cell RNA sequencing approaches allowed us to discover substantial heterogeneity of the lung Tregs during allergic airway inflammation. First, several distinct Treg subsets expressing NLT signature genes were identified. Moreover, the lung Tregs also contained central memory-like Treg subsets, IFN-responsive subsets, and highly proliferative subsets.

While it appeared as if the overall heterogeneity of Treg subsets was comparable between two models of allergic inflammation, re-clustering analysis of the major NLT Treg clusters uncovered that these tissue resident Tregs are strikingly distinct depending on the nature of inflammation. For instance, NLT-like Tregs from alum-induced allergic inflammation were highly enriched with ST2^high^ and CXCR3^low^ cells, whereas NLT-like Tregs from CFA-induced allergic inflammation were ST2^low^ and CXCR3^high^ cells. NLT Tregs also express Lag3 and Gata3, and they appeared to be well separated by the types of inflammation. Indeed, flow cytometric analysis of lung resident vs. intravascular Tregs further confirmed that ST2 expression was primarily observed in lung resident Tregs during alum-induced eosinophilic but not during CFA-induced neutrophilic inflammation.

Positive contribution of the IL-33/ST2 axis to Treg function has previously been investigated. IL-33 induces IL-13 production by ST2^+^ Tregs, which then acquire the ability to control inflammation following lung injury (41). IL-33 was also shown to activate Tregs to suppress IL-17-producing γδ T cells in the lung or in the central nervous system (25, 35). On the other hand, opposite function of IL-33 has also been reported. IL-33 may disrupt Tregs’ ability to control immune tolerance during allergen induced lung inflammation (28). In this study, we found that the lack of ST2 expression in Tregs did not affect their ability to limit allergic inflammation, as T cell production of inflammatory cytokines remained unchanged by the presence of ST2-deficient Tregs. Furthermore, the production of IL-17 by lung γδ T cells was also unaffected in this condition.

Therefore, the IL-33/ST2 axis appears to be dispensable for Treg function to control inflammation in our model. The exact reasons underlying the discrepant findings are not clear. The model allergen or experimental models may play a role. Importantly, ST2^+^ Tregs are abundantly found during alum-induced airway inflammation where IL-33 production is greater than CFA-induced inflammation. It is thus possible that IL-33 produced by lung epithelial cells may be crucial for the generation of ST2^+^ Tregs in the lung. Alternatively, IL-33 may preferentially recruit ST2^+^ Tregs to the site of inflammation. Since ST2^+^ Tregs are rarely found in the lymphoid tissues, ST2 expression may be induced once recruited to the inflamed tissues. Can IL-33 then directly induce ST2 expression in Tregs? IL-33 is able to induce ST2 expression in gastric cancer cells (42). We also observed that IL-33 stimulation of Tregs induces ST2 expression in vitro, although it occurs at a relatively low level (Iamsawat and Min, unpublished observation). Therefore, abundant IL-33 production may be responsible for the ST2 expression of Tregs infiltrating the lung tissue. Whether the IL-33/ST2 axis plays a role in Treg retention in the tissues remains to be determined. Equally plausible is that mediators produced by Th1/Th17 type T cells may inhibit IL-33 production by epithelial cells, from which ST2 expression by Tregs is diminished. IL-33 is also known to stimulate amphiregulin production by Tregs, which is important for tissue repair and remodeling (43). Therefore, IL-33 stimulation in ST2^+^ Tregs may be dispensable for the acute allergic responses while more crucial during chronic phase of the inflammation.

IL-33 is primarily produced by epithelial cells in response to inflammation and tissue stress (44). Unexpectedly, we observed that IL-33 expression in the lung tissue was differentially induced in alum-induced vs. CFA-induced allergic inflammation. Lung epithelial cell production of IL-33 could thus be regulated by IL-33-dependent and - independent mechanism (45). Determining the relationship between epithelial cells’ IL-33 production and effector immunity in response to allergen challenge is a subject of ongoing investigation.

Another important finding from the present study is a trajectory analysis of lung infiltrating Tregs. Pseudotime analysis uncovered interesting cell fate determination that Tregs with highly proliferative and suppressive properties may be subjected to two distinct cellular fates: NLT-like effector Tregs and more central memory-like proliferating Tregs. The former Treg subsets express NLT associated markers and the differentially expressed genes were highly enriched in pathways relevant to immune responses, cytokine stimulation, and T cell activation. On the other hand, the latter Treg subsets express genes associated with ribosomal functions and translation, which are known to be highly enriched in central memory T lineage cells. It is thus possible that highly proliferating (pre-branched) Tregs may represent infiltrating cell types, which then enter the lung tissues and acquire highly suppressive phenotypes to become lung resident Tregs in response to tissue factors including IL-33. One caveat of the single cell RNAseq experiment from this study is that Tregs were FACS sorted from the entire lung tissues without separating lung resident Tregs from intravascular Tregs within the lung. Therefore, it is possible that central memory-like Tregs may be those Tregs that soon leave the lung to re-enter the circulation. Signals involved in the fate determination may be important to investigate. Although the types of lung inflammation showed little impact on cell fate trajectory, re-clustering analysis of NLT-like Tregs highlighted stark differences of Tregs between alum- and CFA-induced inflammation. Lung resident and circulating memory CD4 T cells have been shown to play a distinct role in developing allergic inflammation (31). The precise contribution of lung resident and intravascular Tregs during airway inflammation remains to be investigated.

In conclusion, our results highlight cellular heterogeneity of Tregs in the lung tissue at the peak of allergic airway inflammation. Treg subsets especially those with NLT-like phenotype were distinct depending on the types of the inflammation, i.e., alum-induced eosinophilic and CFA-induced neutrophilic inflammation. ST2^+^ NLT Tregs preferentially emerged in eosinophilic inflammation model. Yet, the lack of ST2 expression in Tregs had little impact on the development of acute inflammation. Future study should test the cellular mechanisms involved in the emergence of different lung resident Treg subsets in response to inflammatory cues.

---

**Figure S1.**
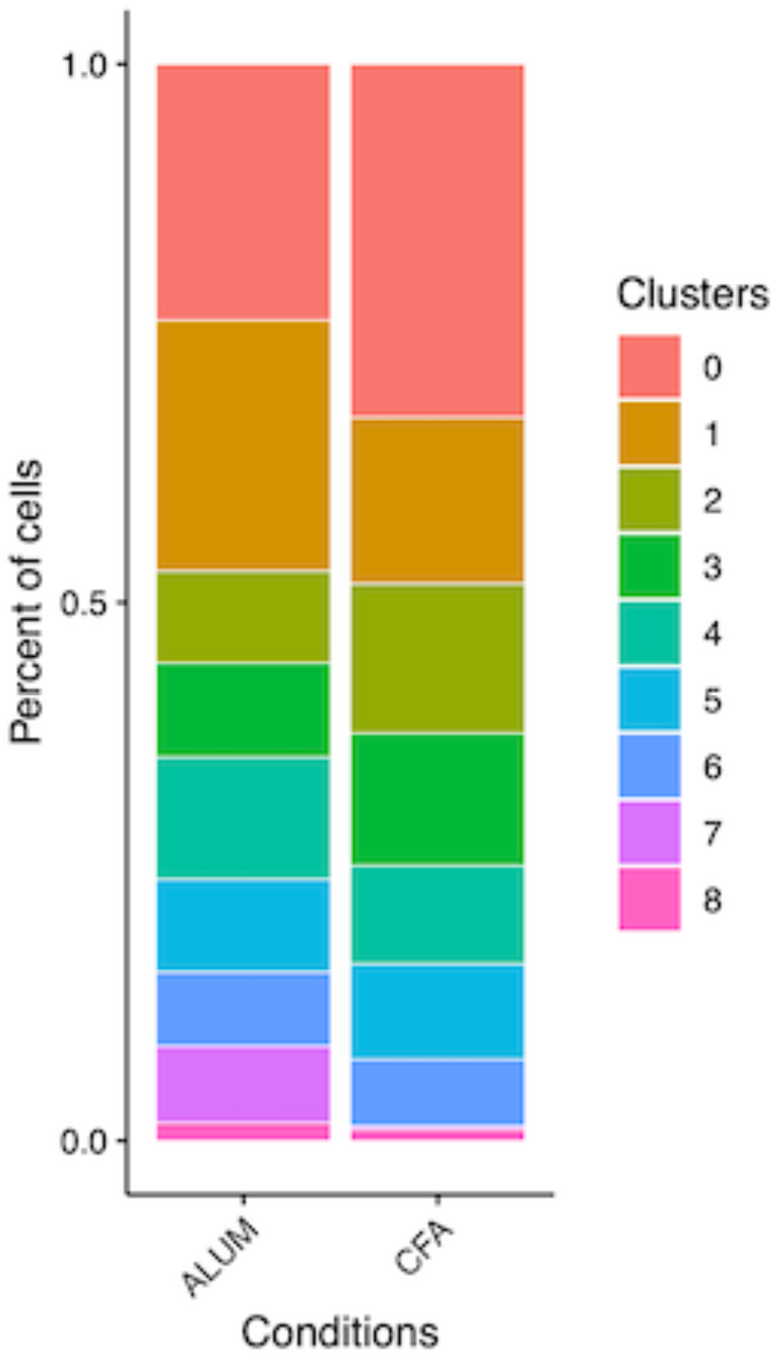
**Proportions of each Treg cluster in alum- and CFA-induced airway inflammation.** Proportions of each cluster of lung infiltrating Tregs from alum- and CFA-induced airway inflammation were calculated.

**Figure S2.**
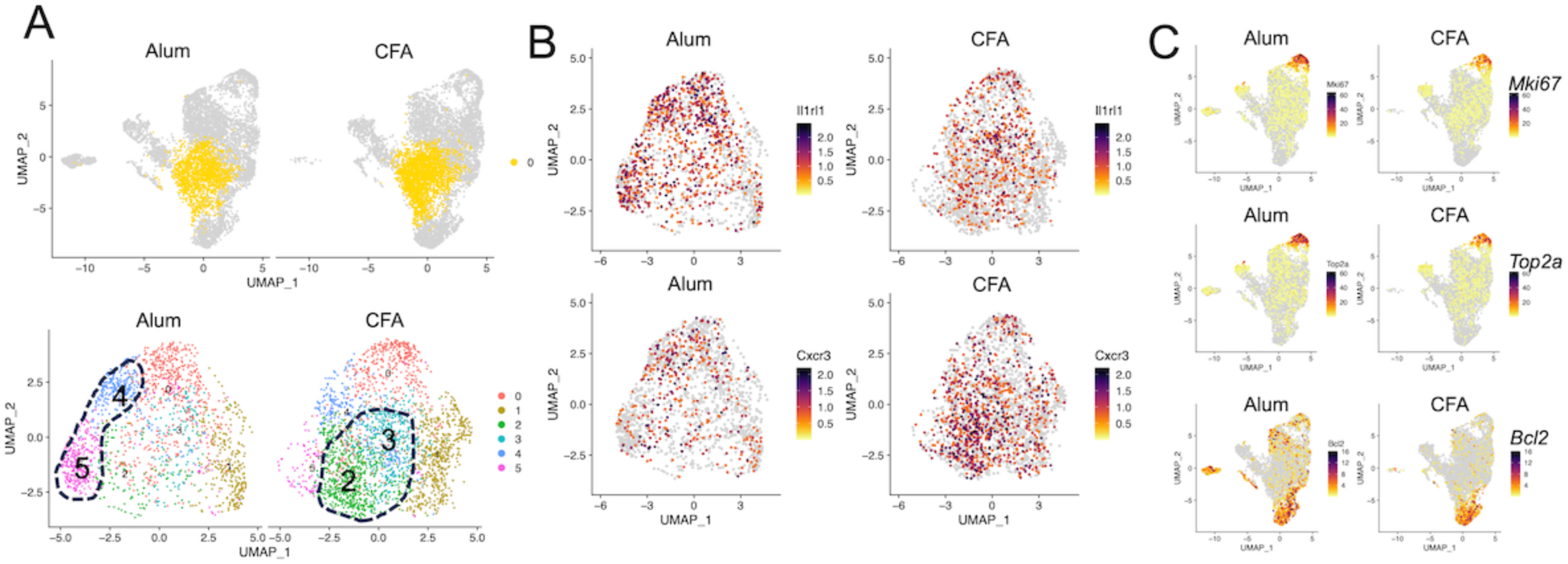
**Re-clustering analysis of NLT-like effector Treg cluster.** (A) Cluster “0” from Figure 2, representative of NLT-like effector Tregs, was re-clustered. UMAP plots show re-clustered Treg populations. (B) Feature plots show *Il1rl1* and *Cxcr3* expressing Tregs. (C) Feature plots showing *Mki67*, *Top2a*, and *Bcl2*-expressing Tregs.

## Notes

This work was supported by NIH grant AI147498 (B.M.)

### Competing Interest Statement

The authors have declared no competing interest.

